# Pre-Stimulus Alpha-Band Phase Gates Afferent Visual Cortex Responses

**DOI:** 10.1101/2021.04.06.438680

**Authors:** Wei Dou, Audrey Morrow, Luca Iemi, Jason Samaha

## Abstract

The neurogenesis of alpha-band (8-13 Hz) activity has been characterized across many different animal experiments. However, the functional role that alpha oscillations play in perception and behavior has largely been attributed to two contrasting hypotheses, with human evidence in favor of either (or both or neither) remaining sparse. On the one hand, alpha generators have been observed in relay sectors of the visual thalamus and are postulated to phasically inhibit afferent visual input in a feedforward manner ^1–4^. On the other hand, evidence also suggests that the direction of influence of alpha activity propagates backwards along the visual hierarchy, reflecting a feedback influence upon the visual cortex ^5–9^. The primary source of human evidence regarding the role of alpha phase in visual processing has been on perceptual reports ^10–16^, which could be modulated either by feedforward or feedback alpha activity. Thus, although these two hypotheses are not mutually exclusive, human evidence clearly supporting either one is lacking. Here, we present human subjects with large, high-contrast visual stimuli that elicit robust C1 event-related potentials (ERP), which peak between 70-80 milliseconds post-stimulus and are thought to reflect afferent primary visual cortex (V1) input^17–20^. We find that the phase of ongoing alpha oscillations modulates the global field power (GFP) of the EEG during this first volley of stimulus processing (the C1 time-window). On the standard assumption^21–23^ that this early activity reflects postsynaptic potentials being relayed to visual cortex from the thalamus, our results suggest that alpha phase gates visual responses during the first feed-forward sweep of processing.

## Results

We quantified afferent visual cortical input in response to task-irrelevant, bilateral, full-contrast radial checkerboard patterns presented in the upper and lower visual field (U/LVF) in separate trials (Figure 1). According to the cruciform model of primary visual cortex, folding patterns in the cortical sheet of V1 (but not V2 or V3) ^17,24,25^ are such that UVF stimuli produce a negative-going C1 ERP and LVF stimuli produce a positive-going C1 (when using an average or mastoid reference) with a scalp focus over central parieto-occipital electrodes ^18,19,26^. Indeed, the stimuli used in this experiment (originally reported in^27^) produced robust C1 responses with polarities, topographies, and timings consistent with striate genesis (Figure 1). The mean amplitude of the C1 component at subject-specific electrodes were 5.26 μV (SEM = 0.61) for LVF and −6.11 μV (SEM = 0.56) for UVF and the average C1 peak latencies were 75.12 ms (SEM = 1.27) and 79.36 ms (SEM = 1.27) for LVF and UVF respectively. Note that bilateral stimulation (likely producing constructive signal interference across left and right V1) gave rise to C1 amplitudes two to three times larger than the typical C1 (cf.^18–20,26^), giving us a strong signal to analyze with respect to the ongoing alpha phase. Following Busch & VanRullen (2010) ^28^, we used the global field power (GFP) during the C1 time window as a metric of afferent V1 input in order to rectify the polarity of the C1 (making larger responses always positive; Figure 1) and because voltage at a single electrode may simply cancel/sum with the phase of ongoing oscillations recorded at that same electrode. For these reasons, GFP has been used in previous research to link pre-stimulus phase to post-stimulus EEG responses, where, for instance, it was found that ~ 7 Hz phase modulates GFP to near-threshold stimuli around 450 ms post-stimulus^28^. As a control, to ensure that alpha phase was not trivially related to post-stimulus GFP in the C1 time-window, we also analyzed a subset of trials (~33%) which contained only a small change at fixation (FIX trials), but no C1-producing stimuli (see Methods).

**Figure 1.**
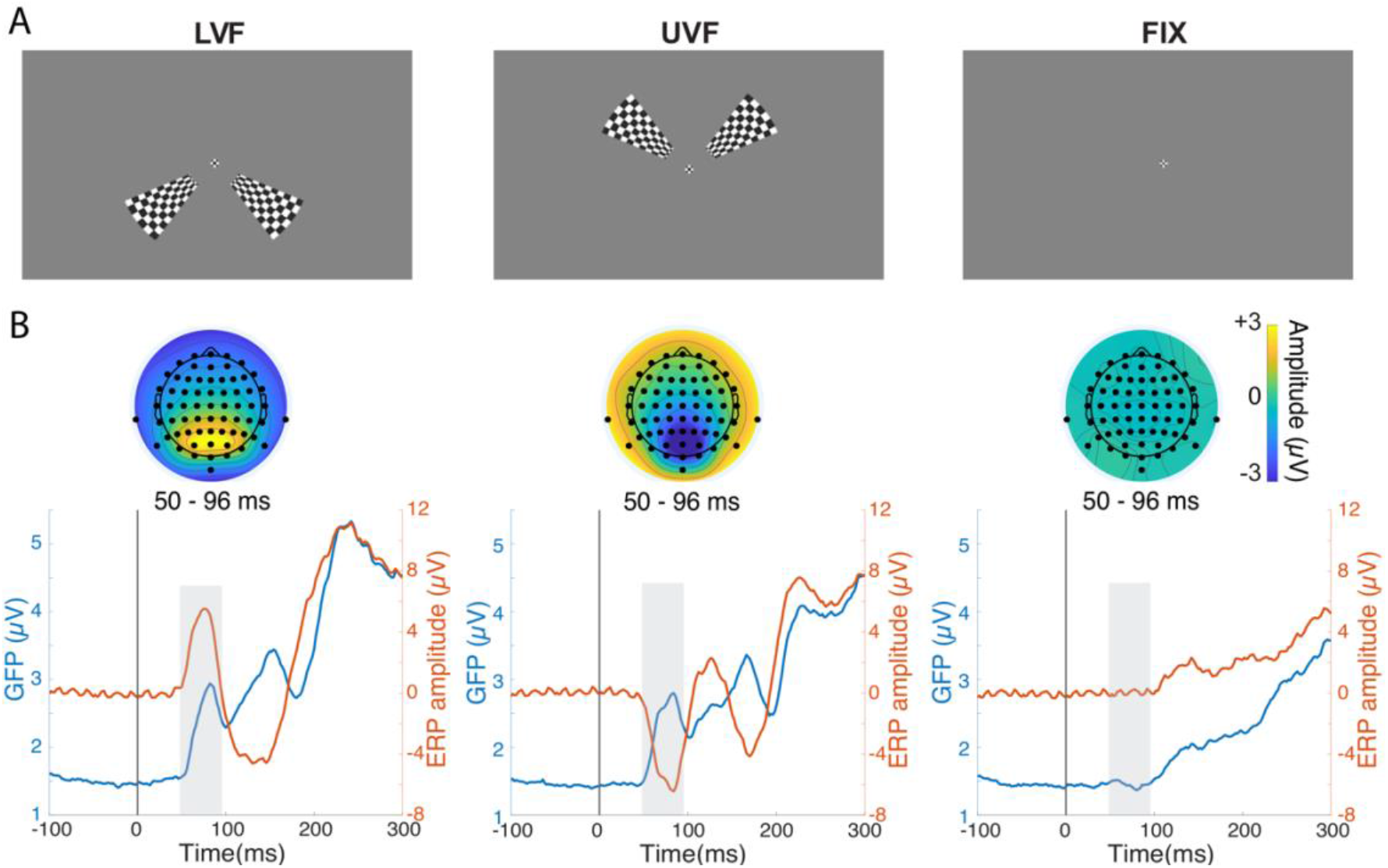
Stimulus Conditions and Stimulus-evoked Responses. (A) The stimuli were task-irrelevant, bilateral, full-contrast checkerboard wedges designed to elicit a robust C1 component. The stimuli were presented in the lower visual field (left column, LVF), upper visual field (middle column, UVF), or were absent (right column, FIX) with equal probability. (B) ERPs (red line) were computed for the subject-specific electrodes with the largest C1 activity. Shaded time windows indicate the time range of the C1 component. The polarity of the C1 reversed across fields of stimulation (LVF vs. UVF) in a manner consistent with striate generation. The C1 was absent on FIX trials. Topographical plots show ERP averaged across 50-96 ms. The GFP (blue line) was calculated by taking the spatial standard deviation of voltage averaged over a 20ms window centered on each participant’s C1 peak within 50-96 ms. GFP provides a global measure of neural response magnitude that also rectifies the polarity reversal of the C1.

We computed circular-linear associations using the weighted inter-trial phase coherence (wITPC)^14,29,30^ to describe the relationship between single-trial GFP amplitudes (averaged over a 20 ms window centered on the C1 peak for each subject) and oscillatory phase across a range of time and frequency points surrounding stimulus onset (extracted via Morlet wavelet convolution). When normalized with respect to the mean and standard deviation of a permuted wITPC distribution, this metric takes on *z* units that reflect the strength of phasic coupling between the linear variable (GFP) and circular variable (phase) relative to a null hypothesis distribution (see Methods). As seen in Figure 2, significant (cluster-corrected, see Methods) wITPC was found between GFP in the C1 time window and pre-stimulus phase for both LVF and UVF trials, but not on FIX trials. For LVF stimuli, GFP was predicted by phase in a cluster of time-frequency points spanning 9 to 18 Hz and −130 to 120 ms from stimulus onset, with a maximum wITPC (z) = 1.74 at 12.35 Hz and 0 ms. For UVF stimuli, GFP was predicted by phase in a cluster spanning 3 to 13 Hz and −150 to 85 ms with a maximum wITPC (z) = 1.65 at 10 Hz and −10 ms. No significant phase-GFP coupling was observed on FIX trials (where only a small fixation change occurred) suggesting that the positive effects on L/UVF trials are not a trivial result of phase predicting GFP at any post-stimulus window but depend on the presence of a C1 component. Scalp topographies from pre-stimulus time points (Figure 2, right panel) indicate an occipital and frontal distribution, consistent with the scalp distribution of the effects of alpha-band phase on visual cortex excitability inferred from TMS-EEG experiments^14,15^.

**Figure 2.**
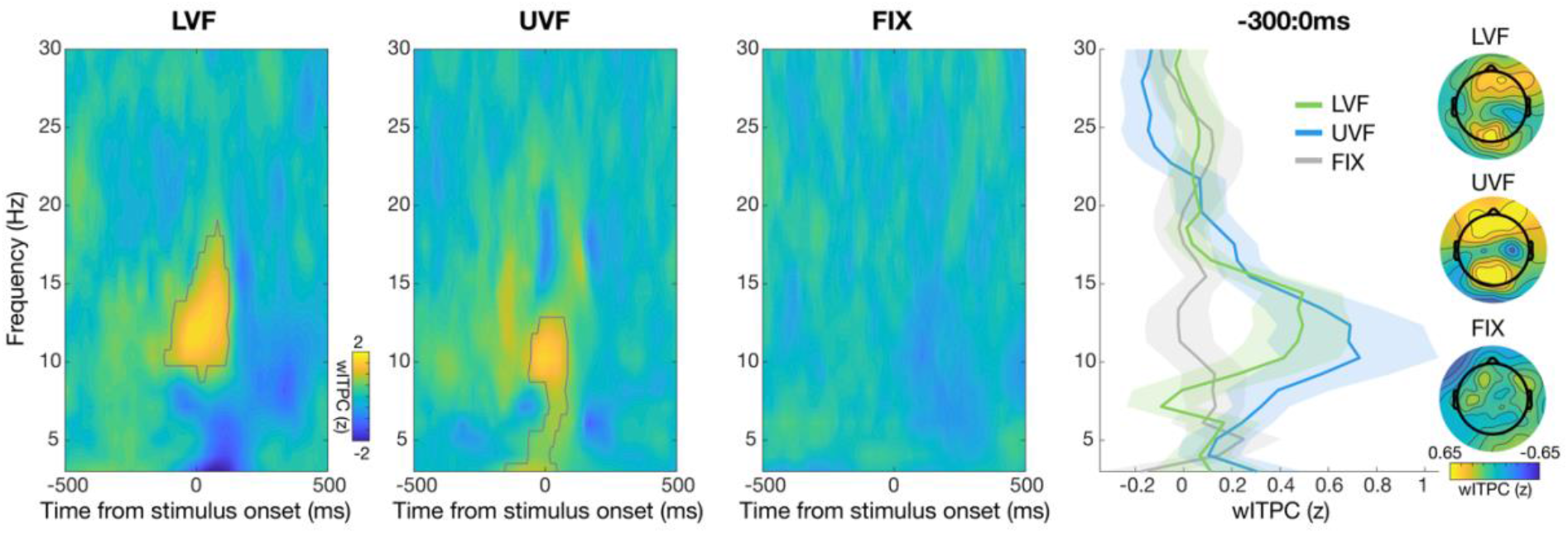
Single-trial time-frequency circular-linear association between phase and GFP. Pseudocolor plots show subject-averaged wITPC(z), which describes the normalized coupling strength between phase across times and frequencies and GPF during each individual’s C1 time-window. The phase-providing electrode was chosen for each individual based on the largest C1 amplitude, but the distribution of wITPC(z) across the scalp from −300 to 0 ms and 8 - 13 Hz is shown in the topographical plots in the right-hand panel. Significant (cluster-corrected) time-frequency points are delimited with a gray line and reveal a modulation of GFP by alpha-band phase (and some adjacent frequencies) just prior to and during stimulus onset for both LVF and UVF stimuli. The effect was absent on FIX trials, when no C1-eliciting stimulus was presented. The right-hand panel shows the wITPC(z) across frequencies for the period from - 300 to 0 ms relative to stimulus onset. A peak in coupling is noticeable in the 8-14 Hz range. Shaded bands indicate ±1 SEM across subjects.

Although a phase effect is expected to be maximal immediately prior to and during stimulus processing and to decay as a function of time before stimulus onset, as has recently been confirmed empirically^10^, it is nevertheless important to confirm that any pre-stimulus phase effect is not confounded by post-stimulus data, which can occur when using a sliding window analysis as done here with wavelets. To this end, we ran an additional analysis using a fast-Fourier transform (FFT) of the 500 ms preceding stimulus onset to exclude any post-stimulus contamination. The FFT approach has the further advantage of allowing us to extract phase at each individual’s peak alpha frequency (see Methods) to complement our mass-univariate wavelet-based approach with a more hypothesis-driven analysis focused on alpha-band oscillations. We sorted post-stimulus GFP in the C1 time window by 7 levels of pre-stimulus alpha-band phase. A 2 × 7 repeated measures ANOVA on post-stimulus GFP with visual field (UVF or LVF) and phase level (1:7, non-shifted) as factors revealed a significant main effect of phase level (*F*(6, 130) = 3.57, *p* < .01). No significant main effect of visual field (*F*(1, 25) = 1.12, *p* = 0.30) or an interaction effect (*F*(12, 300) = .91, *p* = 0.49) were found. We conducted an one-way ANOVA on the GFP on FIX trials with phase level (non-shifted) as within-subjects factor and found no main effect of phase level (*F*(6, 24) = 0.41, *p* = 0.87).

To account for potential individual differences in phase effects^11,16,28^, we re-ran the above analyses after circularly shifting alpha phase for each subject such that the phase with peak GFP became the center phase, which was then removed before statistical testing. We used 2 × 6 repeated measures ANOVA on the GFP, with factors visual field and shifted phase level (excluding the center phase bin used for aligning). The effect of visual field (*F*(1, 25) = 1.14, *p* = 0.30) and the interaction between visual field and shifted phase level (*F*(10, 250) = .60, *p* = 0.70) were not significant. However, as shown in Figure 3, there was a significant main effect of shifted phase level (*F*(5, 125) = 5.41, *p* < .001). For both LVF and UVF, GFP amplitude decreased as pre-stimulus alpha phase deviated from the central phase bin, indicating that pre-stimulus alpha oscillations influenced the afferent sensory response to the U/LVF stimuli. In addition, the effect of shifted phase level on the GFP on FIX trials was not evident (*F*(5, 24) = 0.73, *p* = 0.60) from a one-way ANOVA. This was in line with the results of the wITPC analysis, confirming that the phasic effect relies on the presence of a C1 component. Lastly, we also conducted analyses on the latency of the peak GFP response, which revealed significant modulation (though not of an obviously phasic nature) of alpha phase on GFP peak latency (see Supplemental Materials).

**Figure 3.**
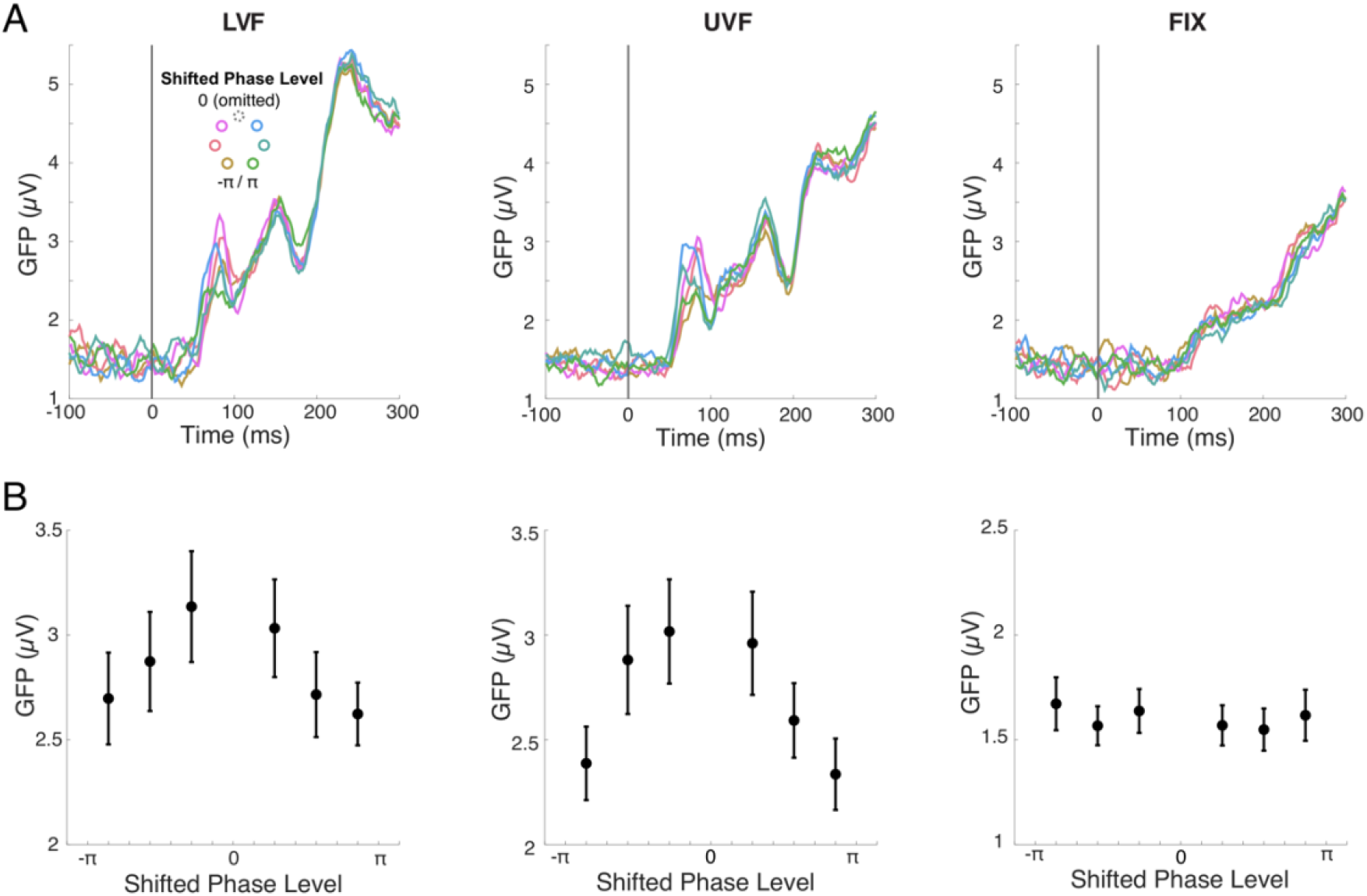
GFP as a function of pre-stimulus alpha phase estimated via FFT. (A) To rule out post-stimulus contamination of phase estimates, single trials were sorted into 7 equally-spaced phase bins between −180° to 180° determined from an FFT using only pre-stimulus data (−500 to 0ms). The phase bins were circularly-shifted to align the phase bin with the largest post-stimulus GFP in the C1 time window to 0°. This central bin was then left out from plotting and statistical tests. Time courses show the pre-stimulus alpha phase modulates GFP for both LVF and UVF trials. No obvious GFP response or modulation by alpha phase was observed within the C1 time window on FIX trials (right). (B) Averaged GFP over a 20ms window centered on each participant’s peak GFP response within the C1 time window decreases monotonically as the shifted phase bins deviate from the removed central bin for both LVF and UVF stimuli. This indicates a phasic effect of pre-stimulus alpha that was absent on FIX trials. Error bars represent ±1 SEM across subjects.

## Discussion

We investigated the influence of alpha oscillations on visual responses by testing the relationship between pre-stimulus alpha phase and post-stimulus GFP measured at the time of the earliest visual-evoked potential in V1, corresponding to the C1 ERP component. According to the “standard model” of EEG genesis, scalp voltage is thought to primarily reflect the post-synaptic potentials of pyramidal neurons^19,21^, meaning the C1 is potentially generated by afferent synaptic input onto V1 from the thalamus. Indeed, a monkey experiment looking at the likely homologue of the C1 component suggests a contribution from layer 4 thalamocortical afferents^31^, although intralaminar synaptic activity within V1 likely also contributes to the C1^32^. Using single-trial circular-linear associations between pre-stimulus phase and post-stimulus GFP, we found a significant effect within the frequency ranges of 9-18 Hz for LVF and 3-13 Hz for UVF, with maximum effects of alpha in both cases. An additional analysis using an FFT ruled out the potential smear of post-stimulus responses into the pre-stimulus window and confirmed the significant modulation of pre-stimulus alpha phase on GFP during the C1 time-window. Analysis of FIX trials, with no C1-producing stimulus, strongly suggest that the effect on U/LVF trials are not a trivial result of pre-and post-stimulus signal autocorrelation. Thus, our results demonstrate that pre-stimulus alpha phase is predictive of the afferent sensory response in the early stages of visual processing

What is the biological mechanism underlying alpha oscillation that accounts for these findings? The visual thalamus, i.e., the lateral geniculate nucleus (LGN) plays a key role in generating the alpha band (8–13 Hz) oscillation measured over visual cortex^1–4,33,34^. Of particular relevance for explaining the current finding is the model of Lőrincz et al^3^, which is based on a combination of in vivo and in vitro recordings of the cat LGN that demonstrated the existence of a specialized subset of LGN neurons that are intrinsically bursting and which synchronize at alpha frequencies via gap junctions^2^. These thalamic bursting neurons were found to drive the local field potential in the LGN and the concurrent scalp EEG at an alpha rhythm that phasically inhibited, via interneurons, the relay mode LGN neurons that carry the afferent visual signals to V1. This model seems well suited to explain the phasic modulation of visual afferent responses in humans that we observed. This implies that the phase of pre-stimulus alpha oscillations could modulate early stages of visual processing by gating the feedforward flow of sensory input from the thalamus and V1. Moreover, a previous analysis of these same data focused on spontaneous alpha amplitude fluctuations found that pre-stimulus alpha amplitude modulates C1 amplitude^27^. Although we take our findings as support for this early feedforward inhibitory model of alpha in humans, we certainly do not intend to downplay the possibility that alpha oscillations also influence the visual cortex from higher- to lower-order cortex in a feedback manner, as has been argued from both monkey and human data^5–8^. These two accounts of alpha are not mutually exclusive.

Several experiments in humans have highlighted a relationship between pre-stimulus alpha phase and different aspects of visual performance in near-threshold detection or discrimination tasks ^11–14,28,35,36^, but see^37,38^. Given the evidence here for a feedforward impact of alpha phase on early visual responses, a reasonable hypothesis is that thalamic gating may underlie the effect of alpha phase on behavior, yet no study has directly linked early sensory responses, alpha phase, and psychophysical performance. Behavior in psychophysical tasks may also be altered by phasic effects of feedback alpha activity, however, sufficient models have not yet been developed that would predict different behavior effects of top-down versus bottom-up alpha effects in visual cortex. Moreover, the precise effect of alpha phase on behavior is itself still not clear, as one report has found effects of alpha phase on response criterion, rather than detection sensitivity ^35^, another paper found effects on discrimination accuracy^13^ (putatively reflecting sensitivity changes), and most other studies have only analyzed hit rates ^10,11,14–16,28^, which are ambiguous between a change in sensitivity or criterion. Moreover, several studies have reported no phase effects on stimulus detection/discrimination^37–39^, though a recent experiment, optimized in many ways to detect an effect, found a rather large change in detection rates of ~20% between stimuli presented during the peak versus trough of occipital alpha^10^. Model-based approaches^40^, as have recently been applied to the study of oscillatory amplitude dynamics^41^, may serve well to better understand the link between oscillatory phase, sensory responses, and perceptual behavior.

## Methods

### The EEG dataset

The EEG dataset was originally collected by Iemi and colleagues^27^ and is available for download at https://osf.io/yn6gb/. In the original study, prior analyses focused on the relationship between pre-stimulus alpha power and C1 responses, but not on alpha phase. The experimental procedure is described in more detail in the original study. In short, 27 participants (mean age: 26.33, SEM = 0.616; 14 females; three left-handed) with normal or corrected vision were presented with a pair of task-irrelevant, full-contrast checkerboard wedges for 100 ms either in the upper (UVF) or lower visual field (LVF) with equal probability. The C1 component can be consistently elicited by checkerboard stimuli on these stimulation trials (see Figure 1). To ensure central fixation, the fixation mark turned into either one of two equally probable targets (‘>’ or ‘<’) for the duration of stimulus presentation. In 33% of trials, only this small central fixation change occurred (FIX trials), giving us a control condition where no C1-eliciting stimulus was presented. The inter-trial interval between stimuli was uniformly selected between 1.8 and 2.4 s. Each subject underwent 810 trials.

The EEG was recorded from 64 electrodes corresponding to the extended International 10–10 system using a Biosemi ActiveTwo system at a sampling rate of 1024 Hz. All channels were referenced online to the CMS-DRL ground electrodes.

One participant did not complete the experiment. We excluded one participant from the analysis because their C1 component was not detected in the LVF after preprocessing. A total of 25 participants were included in the analysis.

### EEG preprocessing

Raw data were preprocessed using custom MATLAB scripts (version R2019b) and the EEGLAB toolbox (Delorme & Makeig, 2004). Continuous recordings were high-pass filtered at 0.1 Hz using a zero-phase Hamming-windowed sinc FIR filter (as implemented in the EEGLAB function *pop_eegfiltnew.m*). EEG data were then downsampled to 500 Hz and segmented into epochs centered on stimulus onset using a time window of −2000 ms to 2000 ms. Individual trials containing eye-blinks and other artifacts during stimulus presentation were inspected visually and removed from data (M = 34.96, SEM = 6.14). Noisy channels were spherically interpolated (M = 0.46, SEM = 0.25) and independent components analysis using the INFOMAX algorithm (EEGLAB function “binica.m”) was used to remove remaining ocular artifacts (M = 1.08, SEM = 0.05). Data were re-referenced offline to the mean of all electrodes and a pre-stimulus baseline of −200 ms to 0 ms was subtracted from each trial.

### Stimulus-evoked responses

To rectify the polarity of the EEG, which is reversed for UVF compared to LVF stimuli (see Figure 1) and provide a global measure of the visual-evoked response during the afferent sweep of cortical activation, we computed the global field power (GFP) of the EEG during the C1 time window. The GFP has been used in prior literature to examine pre-stimulus phase effects on stimulus-evoked responses rather than analyzing the event-related potential (ERP) directly, since the ERP may trivially sum or cancel with the phase of the field oscillations^28^. For the single-trial circular-linear association analysis (wITPC; see below), we computed GFP on single trials by taking the spatial standard deviation of voltage averaged over a 20 ms window centered on each participant’s C1 peak (defined at the electrode with the largest difference between UVF and LVF stimuli in the time-window between 50 and 96 ms post-stimulus; see Figure 1). For the alpha phase binning analysis (FFT; see below), we first computed the ERP for each alpha phase bin and visual field location and then computed the GFP as the spatial standard deviation of each ERP (sometimes referred to as the global mean field power). Then we averaged GFP over a 20 ms window centered on each participant’s peak GFP response within the same time window as the C1 (50-96 ms).

### Single-trial circular-linear association (wITPC)

We took two complementary analysis approaches to study the influence of the pre-stimulus phase on GFP amplitudes during the C1 time window. We first compute the weighted intertrial phase coherence (wITPC)^14,29,30^ as a means of exploring circular-linear associations between phase and GFP across a range of frequencies and time points. wITPC is computed as the resultant vector length, or inter-trial phase clustering (also called the phase-locking factor or inter-trial coherence), of phase angles across trials once the length of each vector has been weighted by the linear variable of interest (here GFP). Note that amplitude information in this analysis is ignored. wITPC was computed separately for each subject, visual field location, time-frequency point, and electrode. Phase was extracted via complex Morlet wavelet convolution at integer frequencies between 3 and 30Hz, with wavelet cycles increasing linearly from 3 to 8 as a function of frequency. A post-wavelet downsampling factor of 5 was applied to speed up subsequent analyses.

Permutation-based statistics were used for both trial-level and group-level analysis. At the trial level, the wITPC value for each condition, time-point, frequency, electrode, and subject was converted to a z-score relative to a null distribution obtained by recomputing wITPC across 1,000 sets of randomly re-ordered trials. Thus, a positive z-score indicates that trials with large GFP values tend to cluster around a specific phase angle more so than would be expected under the same trials but with randomized association. These wITPC (z) values were then tested against zero at the group level using a one-tailed, repeated-measures t-test (since negative wITPC (z) is difficult to interpret), with an alpha of 0.01. Only the electrode with largest C1 for each subject was used in the group level analysis, though Figure 3 displays the topography of average wITPC (z) across all electrodes. For 20/25 subjects, this electrode was POz, 3/25 had PO4, 1/25 had PO3, and 1/25 had Oz. The result of this t-test produced a map of significant time-frequency pixels for each visual field location (i.e., Figure 3). A cluster-based permutation test was conducted to control for the number of comparisons across times and frequencies using the cluster size statistic^30^. On each of 10,000 permutations, a random half the subject’s wITPC (z) values were multiplied by −1 and a one-tailed t-test with alpha set to 0.01 was conducted on this permuted data. The size of the largest significant cluster in the null map was recorded, building a distribution of significant cluster sizes expected under the null hypothesis of no phase-GFP coupling. Only significant clusters in the real data that exceeded the 95th percentile of the distribution of null hypothesis cluster sizes was considered significant. Lastly, to visualize wITPC (z) across frequencies from the pre-stimulus data, wITPC (z) was averaged across the time-window −300 to 0 ms relative to stimulus onset and plotted as a function of the phase-providing frequency (see Figure 3, *right panel*).

### Pre-stimulus alpha-band phase binning (FFT)

To complement the mass-univariate results obtained via wITPC with a more hypothesis-driven analysis of alpha, we conducted a Fast Fourier Transform (FFT) of 500 ms of pre-stimulus data only and sorted GFP in the C1 time window into different bins of pre-stimulus alpha phase. This analysis allowed us to 1) better characterize the effect size of pre-stimulus alpha phase in terms of GFP amplitude modulation, 2) tailor the analysis to each individual’s peak alpha frequency, and 3) rule out any contamination of the results by post-stimulus data since only a pre-stimulus window was extracted for phase sorting.

Pre-stimulus data were extracted from the same set of subject-specific electrodes with the largest C1 response and just from −500 to 0 ms relative to stimulus onset for each trial. Pre-stimulus phase was extracted from single-trials by first linearly detrending each data segment, multiplying the data with a Hamming window, performing an FFT, and extracting the phase angle from the complex Fourier coefficients. Alpha band phase was extracted at each participant’s peak alpha frequency within the range 7 - 14 Hz, taken from the power spectrum of the same pre-stimulus data. Single trials of LVF, UVF and FIX conditions were then separately sorted into 7 equally-spaced bins between −180° to 180°. Following prior work^16,28^, to account for potential individual differences in the specific phase of GFP coupling (related to anatomical differences in how the oscillation field projects to the scalp or differences in V1 conductance latencies), phase bins were circularly-shifted to align the phase bin with the largest GFP amplitude at 0°. This phase bin was then removed from statistical analysis and visualization to remove bias. Statistical tests are reported for both the shifted and non-shifted data (see Results).

## Supplemental Information

### Supplementary Results

In the analysis using an FFT of the 500ms preceding stimulus onset, we sorted post-stimulus GFP in the C1 time window by 7 levels of pre-stimulus alpha-band phase. In addition to the statistical analysis on the GFP amplitude, we conducted a 2 × 7 repeated-measures ANOVA on the peak latency of the GFP to test the phasic modulation in the latency of visual responses with one factor being visual field (UVF or LVF) and the other factor being phase level. We found no significant main effects of visual field (*F*(1, 25) = 0.61, *p* = 0.44) or non-shifted phase level (*F*(6, 130) = 0.62, *p* = 0.71) and no significant interaction effect (*F*(12, 300) = 1, *p* = 0.38). A one-way ANOVA on the GFP on FIX trials with phase level as the within-subjects factor showed no main effect of non-shifted phase level (*F*(6, 24) = 0.88, *p* = 0.51). We then circularly shifted alpha phase for each subject such that the phase with peak GFP became the center phase bin and removed this center bin from any further statistical analysis, as was done for the analysis on GFP amplitudes. A 2 × 6 repeated measures ANOVA on the peak latency of GFP, with visual field shifted phase level as factors, revealed that shifted phase level significantly influenced the peak latency of GFP (*F*(5, 125) = 3.99, *p* < 0.05), whereas visual field did not (*F*(1, 25) = 0.30, *p* = 0.59). The interaction between visual field and shifted phase level on the peak latency was significant (*F*(10, 250) = 2.38, *p* < 0.05). We did not find any difference in the peak latency between shifted levels on FIX trials (*F*(5, 24) = 1.74, *p* = 0.13). Although the statistical effects indicate some impact of alpha phase (only once shifted) on peak GFP latency, the pattern driving these effects, as seen in Supplementary Figure 1, does not obviously appear phasic (e.g., decreasing/increasing with deviation from the central phase bin), and is not clearly consistent for UVF and LVF stimuli, in contrast to the effects on GFP amplitude reported in the main paper. We tentatively conclude that there may be effects on visual response latency, but not in a phasic manner.

**Supplementary Figure 1.**
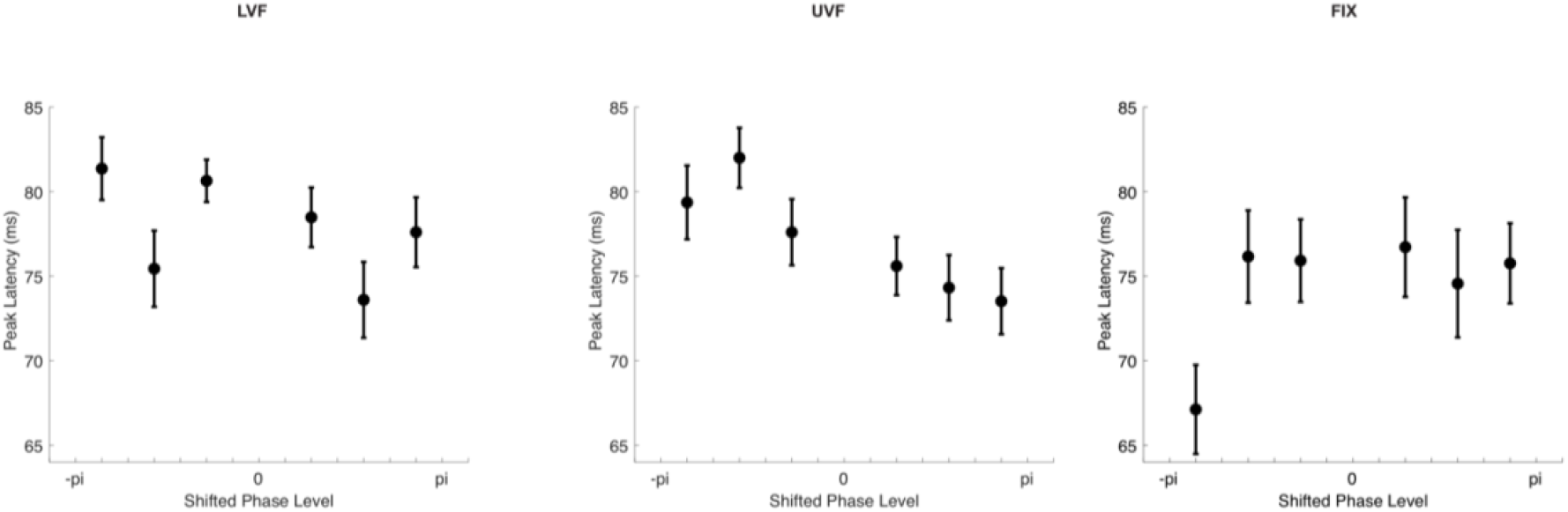
Peak latency of GFP centered on each participant’s peak GFP response within the C1 time window changed on different phase bins, but the patterns for LVF and UVF were dissimilar and neither was obviously phasic in nature. No statistical effect was observed on FIX trials. Error bars represent ±1 SEM across subjects.

